# MetaPhage: an automated pipeline for analyzing, annotating, and classifying bacteriophages in metagenomics sequencing data

**DOI:** 10.1101/2022.04.17.488583

**Authors:** Mattia Pandolfo, Andrea Telatin, Gioele Lazzari, Evelien M. Adriaenssens, Nicola Vitulo

## Abstract

In the last decades, a great interest has emerged in the study and characterisation of the microbiota, especially the human gut microbiota, demonstrating that commensal microorganisms play a pivotal role in normal anatomical development and physiological function of the human body. To better understand the complex bacterial dynamics that characterize different environments, bacteriophage predation and gene transfer need to be considered as well, as they are important factors that may contribute to controlling the density, diversity, and network interactions among bacterial communities. To date, a variety of bacteriophage identification tools have been developed, differing on phage mining strategies, input files requested and results produced; however, new users approaching the bacteriophage analysis might struggle in untangling the variety of methods and comparing the different results produced. Here we present MetaPhage, a comprehensive reads-to-report pipeline that streamlines the use of multiple miners and generates an exhaustive report to both summarize and visualize the key findings and to enable further exploration of specific results with interactive filterable tables. The pipeline is implemented in Nextflow, a widely adopted workflow manager, that enables an optimized parallelization of the tasks on different premises, from local server to the cloud, and ensures reproducible results using containerized packages. MetaPhage is designed to allow scalability, reproducibility and to be easily expanded with new miners and methods, in a field that is constantly expanding. MetaPhage is freely available under a GPL-3.0 license at https://github.com/MattiaPandolfoVR/MetaPhage.

## Introduction

Bacteriophages, viruses infecting bacteria, are increasingly studied as part of the resident microbiota in any environment, from the global oceans over soils to the human gut. Where early studies used BLAST-based approaches and read profiling to describe the virome based on its relationship to genomes in reference databases, the current gold standard includes the reconstruction of (near) complete viral genomes through assembly and viral contig identification. The latter approach has resulted in the IMG/VR v3 database containing 2,332,702 distinct viral genomes or UViGs (uncultivated virus genomes) (Roux et al., 2021), or a number of human gut-associated virus databases, e.g. the Gut Virome Database (Gregory et al., 2020) and the Gut Phage Database (Camarillo-Guerrero et al., 2021). Beyond assembly and identification of viral contigs, clustering in viral Operational Taxonomic Units (vOTUs) that are aligned with the phage species level has emerged as the gold standard to reduce dataset complexity (Roux et al., 2019; Turner et al., 2021). Using 95% average nucleotide identity over 85% of the aligned fraction, the IMG/VR database contains 933,352 vOTUs. The advent of high-throughput sequencing and metagenomics led to the direct analysis and identification of the genetic material in environmental samples, overcoming the barrier of culturability. Two different approaches are used nowaday; sequencing of the whole metagenome followed by computational viral sequences isolation or the physical separation of viral fractions to produce the so-called metavirome.

The general strategy to identify and classify viruses from metagenomics datasets usually involves five steps: (i) pre-processing of the raw reads; (ii) filtering of non-targeted reads; (iii) assembly of short sequence reads; (iv) virus identification and classification and (v) post-process of the viral contigs sequences (Nooij et al., 2018). Several tools exist to perform each step, but while pre-processing, filtering and assembly can rely on gold standard softwares and workflows, viral identification and classification may vary greatly between different tools.

Many of the virus mining tools were implemented in the last decade, and differ on the approach used, such as prophage detection, homology, machine learning, random forest or hybrid approaches, which is a combination of some of the aforementioned (Ho et al., 2021).

Furthermore, one major drawback of virus metagenomic analysis is the variety of tools used in each step, which needs to be installed separately, may require different data formats and an extended set of dependencies, which may clash in their versions. In addition, the output formats from one set of tools may not be compatible with downstream programmes.

To assist the non-specialist in the decision-making process and facilitate workflow management, we present here MetaPhage, a fully automated computational pipeline for phage detection, classification and quantification of metagenomics data. The pipeline offers the possibility to skip some of the steps or recover the analysis in case of execution errors. To guarantee scalability and reproducibility, Metaphage was implemented in Nextflow (NF) (Di Tommaso et al., 2017) [v21.04.0], a workflow manager that uses software containers to allow an easy installation. The pipeline can be run on a single computer or parallelized on an HPC cluster. Metaphage also implements a novel algorithm that allows an automatic taxonomy classification of the viral contigs from the vConTACT2 network graph. Results for each step of the analysis are reported on a rich and easy-to-read HTML report that can be opened and inspected on any web browser.

## Materials and Methods

### Overview

The pipeline was implemented in Nextflow (NF) [v21.04.0], a portable, scalable, and parallelizable workflow manager. MetaPhage (MP) includes four different phage-mining tools to identify the viral sequences in prokaryotic metagenome data, together with thirteen other tools for quality control, assembly, clustering, annotation and classification, as shown in **Fig. 2**. These tools are integrated into MP using Singularity [v3.7.1] (Kurtzer et al., 2017), a container platform that allows to package software in a single file (i.e. container), which is portable and reproducible; this approach frees the user from the burden of installing software and dependencies, which may be difficult to beginners in bioinformatics.

**Fig 1.**
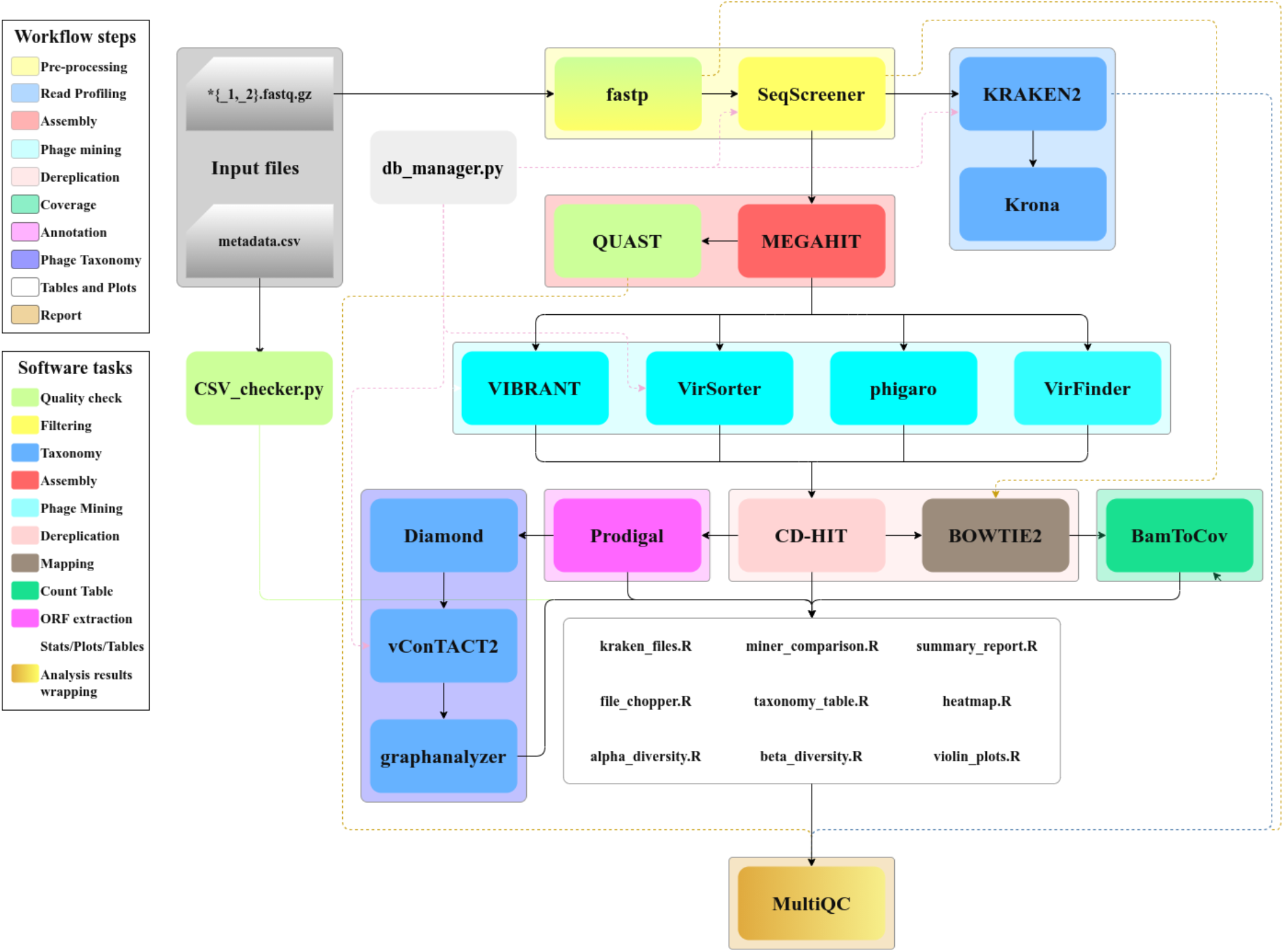
MetaPhage workflow DAG chart. Input sequences can either be SE or PE, while metadata are .csv files with a strict format style, as described in the MetaPhage manual. By configuration file or command line, the user can decide to skip several steps or phage identification tools.

**Fig 2.**
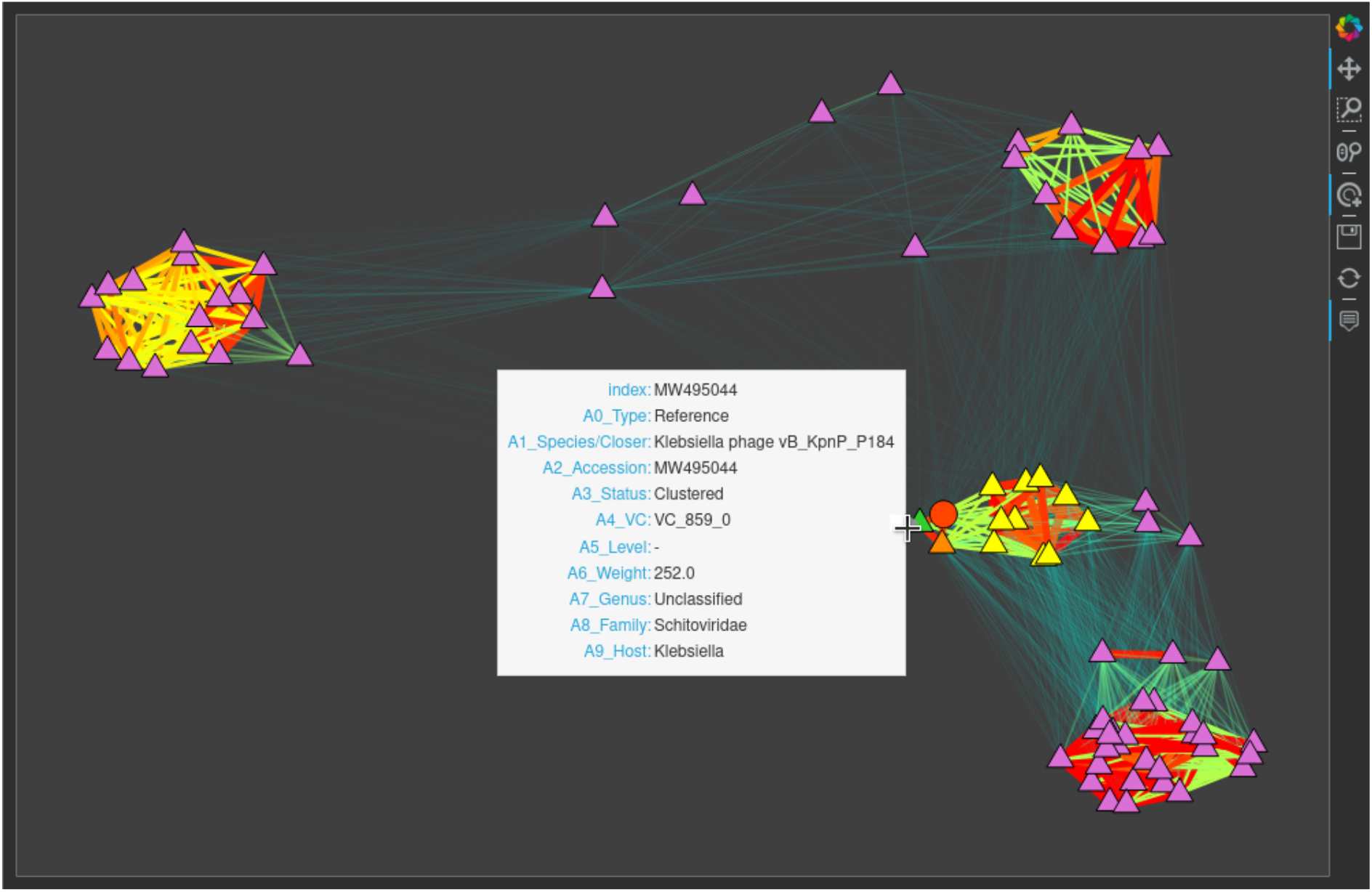
Interactive subgraph produced by graphanalyzer for a VCS. Triangles and circles represent reference genomes and VCS, respectively. This VCS and the reference genome from which it inherits the taxonomy are depicted in red and green, respectively. Orange nodes are sub-clustered together with this VCS, while yellow nodes are only clustered together and belong to a different sub-cluster. The width and color of edges are proportional to the similarity between two nodes, from thin transparent aquamarine (weaker similarity) to thick opaque red (strongest similarity) shading from green, yellow and orange in-between. Nodes are positioned approximately respecting the similarity between them, and this tends to make clusters visible at first sight. When opened with a browser, this subgraph is interactive: the user can zoom and drag it, and can hover a node with the mouse to show its properties, like ‘species’, ‘genus’, ‘family’, ‘viral cluster’ and more.

Therefore, MP offers an installation-free, dependency-free and conflict-free workflow execution, regardless of the operating system.

The six mandatory databases needed for the analysis are automatically downloaded and stored by the pipeline on the first execution; moreover, users can define different, custom databases through the MP config file. Almost every pipeline step can be skipped, and many options are fully customizable by the user through the MP config file; if no custom config is provided, the pipeline runs with default options.

The details of the main steps are reported here below.

### Database download

MP relies on several external databases: (i) *Phix*, used to remove the calibration control of sequencing runs during the *pre-processing* step; (ii)

*MiniKraken2_V1* as default Kraken2 database used in the *read classification* step (available here: https://benlangmead.github.io/aws-indexes/k2); (iii) VIBRANT (Kieft et al., 2020), (iv) Phigaro (Starikova et al., 2020) and (v) VirSorter (Roux et al., 2015) databases containing all the dependencies needed by the relative tools; (vi) INPHARED (Cook et al., 2021) database which provides a collection of curated bacteriophage genomes from GenBank and related metrics, together with several input files useful in bioinformatic pipelines.

### Pre-processing and sequence manipulation

MP takes in input single-end/paired-end Illumina fastq files; at first, the fastp program [v0.20.1] (Chen et al., 2018) is used to remove user defined adapter sequences (TrueSeq adapters by default) and to trim raw reads using a sliding window approach. The *-5* and *-3* options are used to move from 5’ to tail and from 3’ to head of each sequence, trimming where quality falls below a user-defined threshold set with the *--cut_mean_quality* option (15 by default). If no adapter sequences are provided (via *--adapter_sequence* and *--adapter_sequence_R2* fastp options respectively), the tool tries to automatically detect the correct adapters for the input sequences. The *SeqScreener* module from HTStream package [v1.3.3] (*s4hts/HTStream*, 2021) is then run, to discard contaminants mapping the *phiX174* genome (default) or any user defined genome, and the resulting phix-depleted and pre-processed reads are sent to the next steps, which run in parallel: *Read Classification* and *Assembly*.

Manipulation of FASTA files is performed via SeqFu [v1.9.6] (Telatin et al., 2021).

### Read classification

Microbial taxonomy classification is performed by Kraken2 tool [v2.1.2] (Wood et al., 2019). In order to help the user’s further downstream analysis of the microbial classification, Kraken2 is run with the option *–report-zero-counts*, which displays all taxa, even those not assigned to any reads. The Kraken2 output is then passed to Krona [v2.8] (Ondov et al., 2011), which summarizes the results creating interactive html pie-charts. This step provides a general overview of the microbial composition within the samples, including any potential contaminants. Notice that it is not tailored for the identification of phages, but to provide an ecological context that may be useful for a better interpretation of the results.

### Genome Assembly

Reads are assembled using the MegaHIT assembler [v1.2.9] (Li et al., 2015) with default parameters. The assembled scaffolds are evaluated on their quality through the *Assembly quality check* step; this is performed by metaQuast [v5.0.2] (Mikheenko et al., 2016), a tool which evaluates and compares metagenome assemblies based on alignments to close references provided by the user. Since reference genomes are not provided, the tool uses BLASTN to identify the metagenome content, aligning contigs to SILVA 16s rRNA database. To enable ribosomal RNA gene finding, the option *--rna-finding* is used. The resulting report includes several per assembly statistics, such as contigs cumulative length, N50 and GC content, which are plotted in an html report file, with interactive graphs. Concurrently, the assembled scaffolds outputted by MegaHIT are passed to the *Phage mining* step, for the identification of the bacteriophage genomes.

### Phage mining

MP exploits four different phage mining tools which run in parallel to identify viral sequences in the provided contigs/scaffolds: VIBRANT [v1.2.0], Phigaro [v2.3.0], VirSorter [v1.0.6] and a parallelized version of VirFinder [v1.1]. The first three tools can be run with the default database available with the program or with a user provided database. VIBRANT runs with default parameters (minimum scaffold length requirement set at 1000 bp, minimum number of open reading frames (ORFs) per scaffold set at 4). The option *-virome*, which increases sensitivity for virome datasets, removing non-viral scaffolds, is used when the MP option *–virome-dataset* is set to true by the user. Since GC quantity levels may vary depending on the dataset of input, Phigaro runs in basic mode (see Starikova et al., 2020 for details). VirSorter and VirFinder run with default parameters. The identified viral contig sequences are then sent to the *Dereplication* step.

### Dereplication and viral contigs comparison

The dereplication process consists of clustering viral contigs according to a nucleotide similarity threshold. Viral contigs are dereplicated using CD-HIT [v4.8.1] (Fu et al., 2012) with sequence identity threshold=0.95, word size=9, and alignment coverage=0.85. This step makes use of the “accurate mode “ (*-g*: 1) by which sequences are clustered to the most similar cluster that meets the threshold. In order to compare the results of the different phage-mining tools, an R script based on the UpSet R library [v1.4.0] (Conway et al., 2017) generates a comparison plot as shown in **Fig. 3, panel A**. This graph provides a general overview of how many viral contig sequences (VCS) are identified by each miner (i.e. after dereplication), and how many are mined by two or more tools. VCSs are the input for the following concurrent steps of the pipeline: *Mapping* and *Annotation*.

**Figure 3.**
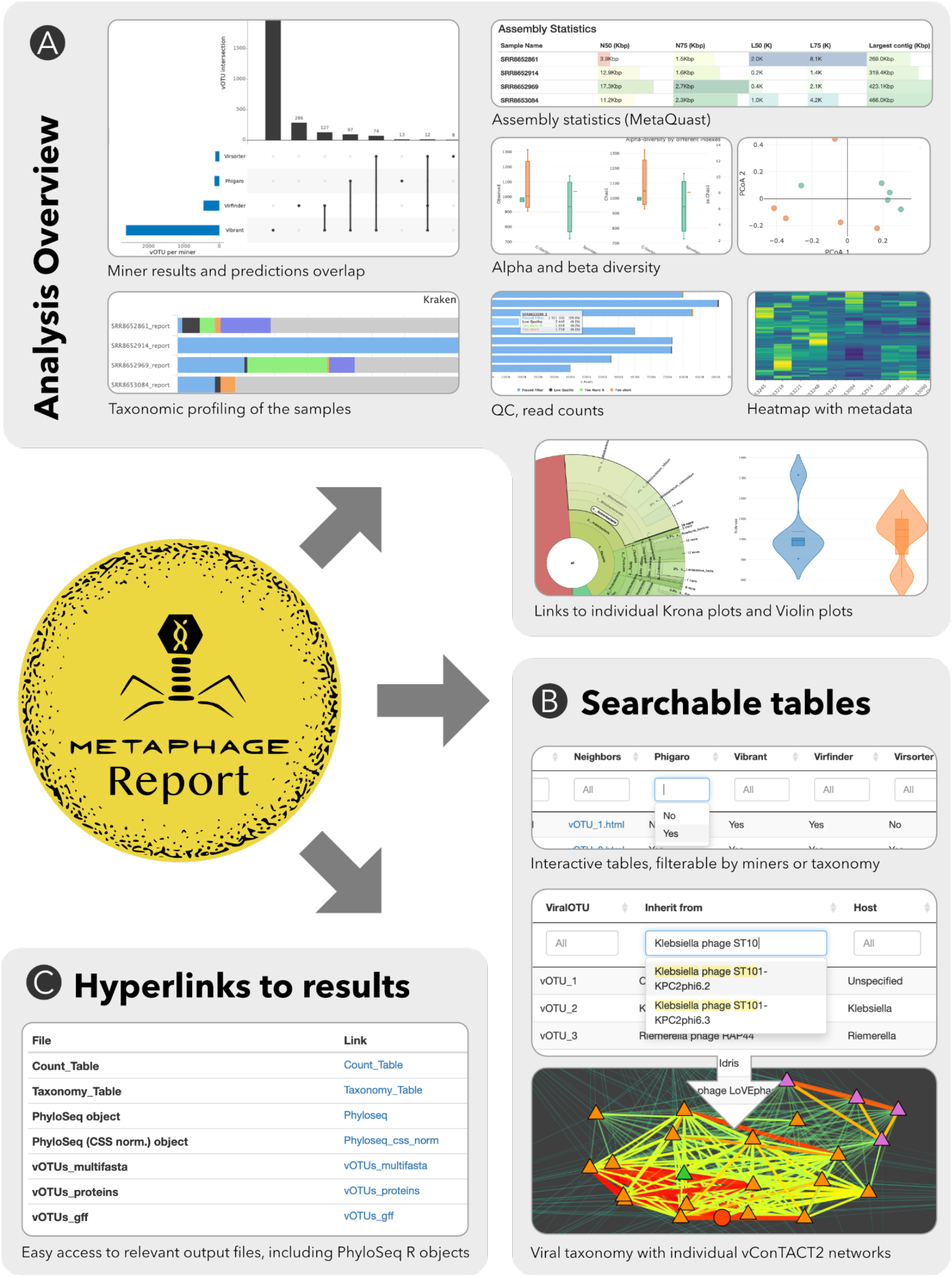
MetaPhage produces an HTML report with multiple sections, which can be divided in three main categories: analysis overview, searchable tables and hyperlinks to results. **(a)** Analysis overview includes panes to inspect the overall quality of the reads (*fastp*), and taxonomical composition of the whole sample (*Kraken2*) including interactive plots for each sample (*Krona*), the assembly metrics (*metaQuast*), plus custom plots specific to the viral diversity and produced with R by the pipeline (alpha and beta diversity, heatmap, violin plots). **(b)** The searchable tables include a summary of the taxonomic analysis of the viral OTUs (vOTUs) as performed (vConTACT2), and custom filters can be added to exclude some miners or to restrict the search to specific phage clusters in the network. For each vOTU, a link to the individual network (as performed by the graphanalyzer script). **(c)** A dedicated section reports links to the main files produced by the pipeline for downstream analyses, including the raw counts table, the taxonomy table and PhyloSeq objects for downstream analyses in R.

### Mapping and count table construction

Bowtie2 [v2.4.2] (Langmead and Salzberg, 2012) (Langmead et al., 2019) is used to quantify VCSs in each phix-depleted, pre-processed reads, mapping them back and processing the results with SAMtools [v1.11] (Li et al., 2009). The resulting alignment is sent to the *Count table construction* step, which makes use of the *bamcountrefs* module (default parameters) of the BamToCov tool [v2.0.4] (Telatin and Birolo, 2022) in order to create a count table. The module is run with option *–multiqc* to output the table in the correct format, and the resulting table presents the raw reads count, with no normalization applied. The table is further modified by a python script in order to change the VCSs label in vOTUs, then is passed to the *Plots&Report* step.

### Annotation and phage taxonomy

Concurrently, Prodigal [v2.6.3] (Hyatt et al., 2010), a prokaryotic and viral gene recognition tool, is used in metagenome mode to predict viral ORFs from the VCSs. This tool makes use of an unsupervised machine learning algorithm that automatically learns the genome properties (start codon, ribosomal binding site motifs, coding statistics), without a training data set. The metagenome mode option (*-p meta*) is mandatory when working with mixed samples, such as the assembly of a metagenome.

The resulting protein sequences are then passed to the viral taxonomy step, which uses vConTACT2 [v0.9.19] (Jang et al., 2019) a tool that performs network-based clustering of viral sequences and specifically designed to work on metagenomics data. vConTACT2 uses whole-genome gene-sharing profiles between a reference database of curated and annotated bacteriophages and the user viral sequences, to build networks for phage taxonomy. One of the key steps in the vConTACT2 pipeline is an all-versus-all predicted protein comparison that is used to generate clusters of protein families. The reference database and the user viral proteins are merged in a single file to perform an all-versus-all similarity search. Depending on the size of the reference database, this step can require a long time, as suggested by the authors of the tool; moreover, the network is rebuilt every time new data are added, extending, even more, the time consumption (Jang et al., 2019).

We reason that, if the reference database does not change, we can perform the all-versus-all protein comparison of the reference just once, as the results will not change. In order to try to speed up the process, we decided to split the all-versus-all search of the reference database from the user viral proteins search, running vConTACT2 using *vConTACT2_proteins*.*faa and vConTACT2_gene_to_genome*.*csv*, two files contained in the INPHARED database. For a detailed description of the approach, see the *Custom vConTACT2 script* paragraph in the Supplementary.

The similarity network and the clusters computed by vConTACT2 are then processed by graphanalyzer [v1.2.1], a novel script we developed to automatically inherit the taxonomy of each VCS from its closest reference genome. This reference is chosen by looking for connected genomes inside the network, and prioritizing those included in the same cluster. This saves much time and effort, since vConTACT2 does not classify a VCS directly but instead would leave the classification up to the researcher’s extensive manual inspection. Moreover, graphanalyzer produces a summarizing taxonomy table (.csv file) which is passed to the *Plots&Report* step, as well as an interactive subgraph for each VCS that helps the user to explore its taxonomic context. The tool was written in Python and is freely available at https://github.com/lazzarigioele/graphanalyzer. Further details of its algorithm are discussed in the Results section.

### Plots and Report generation

In order to produce interactive plots and tables, several R scripts make use of the vOTUs count table produced by BamToCov, the taxonomy table produced by *graphanalyzer* and the opportunely formatted metadata (.csv) file (see Results: MetaPhage output section for in-detail explanation). These scripts make use of different R libraries, such as plotly, Phyloseq (McMurdie and Holmes, 2013), metagenomeSeq (Paulson et al., 2013), tidyverse (Wickham et al., 2019), seqinr (Charif and Lobry, 2007), heatmaply (Galili et al., 2018), DT (Xie et al., 2022), Biostrings (Pagès et al., 2022). The pipeline steps are summarized via MultiQC (Ewels et al., 2016), a software which produces a comprehensive interactive report, available via web browser. This report integrates several software output reports (fastp, Kraken2, metaQuast), together with the custom interactive plots, as shown in **Figure 1**.

## Results

### Overview of the MetaPhage workflows

Metaphage implements a fully automated computational pipeline for phage detection, classification and quantification of metagenomics data (Fig 1). Several parameters are available to customize the analysis workflow, while the pipeline offers the possibility to skip some of the steps or recover the analysis in case of execution errors.

The minimal input consists of the metagenomics reads in FASTQ format. Optionally, but recommended, the user can also provide a metadata file containing information about the samples that can be used to customize the output. The pipeline is implemented in Nextflow (NF) [v21.04.0], a scalable and reproducible scientific workflow manager that uses software containers to allow an easy installation. A fully documented installation and usage guide is available at https://mattiapandolfovr.github.io/MetaPhage/. The workflow can be run on a single computer or on a HPC taking advantage of different job schedulers such as Slurm, Torque or PBS. MP also implements a novel algorithm that allows an automatic taxonomy classification of the viral contigs using vConTACT2 clusters. Results for each step of the analysis are reported on an easy-to-read html report that can be opened and inspected on any web browser.

### Dependencies

MetaPhage dependencies are available from the BioConda project (Grüning et al., 2018), and can be exposed to the pipeline as a Conda environment, as a Docker container or as a Singularity image, all based upon a predefined environment (YAML) distributed with the pipeline.

### Viral taxonomy classification

vConTACT2 is the taxonomy classification tool implemented in MP. Briefly, it uses viral genomes as nodes of a monopartite network from which it derives viral clusters. Genomes are grouped based on protein content and similarity, and the resulting clusters are largely concordant with the ICTV viral taxa (Jang et al., 2019).

vConTACT2 tries to split clusters into subclusters, where clusters are a rough approximation of phage (sub)families, and subclusters are the same for genera. For each sequence in the network, vConTACT2 assigns a label representing the clustering “status”. A user can encounter several status descriptions, like “Clustered”, when the sequence clearly falls in a defined cluster and subcluster. More complex status classifications are also possible, like “Overlap” when a sequence fits two or more clusters equally well, “Singleton’’ when a cluster is composed by that lonely sequence, or “Outlier” when the sequence is not assigned to any cluster.

The network produced by vConTACT2 has viral sequences (reference genomes and VCS) as nodes and its edges are weighted proportionally with the similarity between two nodes. This network can be inspected using Cytoscape, and this helps the user to deduce the taxonomic context of each VCS, including the most difficult cases. Unfortunately, repeating this manual inspection for each VCS is very time-consuming. To try to automate the process, we developed a novel Python script named *graphanalyzer*, that assigns an optimal taxonomy to every VCS following the same process a user would do manually.

For each VCS to be classified, graphanalyzer searches the corresponding node in the similarity network, and retrieves the list of nodes directly connected to it (its “neighbors”), that are also clustered in the same cluster of the VCS (if any). The list is then ordered by the weight of the edge connecting the node to the VCS, and the taxonomy is inherited from the first reference genome found in this sorted list. If no reference genome is found, graphanalyzer retrieves a new sorted list with all nodes that are simply neighbors, regardless of the clustering information. Several iterations of the algorithm are performed until there’s no new taxonomic assignment. At every new iteration, previously classified VCSs are used as reference genomes.

In this process, the taxonomic classification for each VCS can be more or less confident depending on the number of iterations done and the clustering status. To be more conservative, we decided to stop the inherited taxonomy at the level of subfamily if the status is “Singleton” or “Overlap”, and at the level of order if the taxonomy was inherited from a genome not included in the same cluster. Otherwise, the taxonomy is kept until the level of genus.

For each VCS, graphanalyzer also produces an interactive subnetwork that can be explored to retrieve extra information without the need for Cytoscape (Fig. 2). Further information on graphanalyzer including its installation and usage, a description of its main algorithm and its classification levels, and a detailed user guide to the interpretation of the taxonomy table and the interactive subnetworks it produces, are freely available in its public repository https://github.com/lazzarigioele/graphanalyzer.

### MetaPhage report

MetaPage results are summarized in a rich, easy-to-read HTML report that can be opened in any web browser. The report is organized in different sections for each step of the workflow.

The *Miner Comparison* section shows the number of VCSs identified by the different phage mining tools and allows a direct comparison of the performance of each software (**Fig 3, panel A**). The *Summary Table* section displays per-sample general statistics, such as vOTUs total count, abundance, vOTUs length (minimum, average and maximum) and vOTUs distribution in a searchable table. The *Taxonomy Table* section (**Fig 3, panel B**) is an interactive and searchable per-vOTU table, displaying taxonomy information and host association information (retrieved using the information contained in the INPHARED database). The taxonomy is automatically assigned using *graphanalyzer* tool, and it stops at the genus level, if available. For each vOTU it is also possible to visualize the corresponding network generated by *graphanalyzer*, together with its nucleotide consensus sequence, predicted protein sequences (FASTA format) and coordinates files (GFF3 format). The table allows the user to query for any combination of phage miners to restrict the search to only the vOTU of interest. The *vOTUs Distribution* (**Fig 3, panel A**) section contains the links to the generated heatmap and violin plots, both interactive plots which give information about the vOTUs abundance (log count) and distribution across the samples.

The report also provides alpha and beta diversity metrics in the omonim sections, to describe the within-sample complexity and between-samples diversity. In particular, the alpha diversity is calculated using different metrics such as Observed, Shannon, ACE, Simpson and Fisher, while the beta diversity is calculated using the Bray-Curtis and Jaccard indices.

Several important files generated during the analysis and useful for downstream analysis are also directly linked within the report, for easy and fast access, in the *Interesting Files* section (**Fig 3, panel C**). In particular, the vOTU count table that reports the abundance of each vOTU in each sample, the taxonomy table, multi FASTA files containing the vOTUs nucleotide and protein sequences and the coordinate of the predicted proteins on the corresponding contigs in gff3 format. Moreover, a PhyloSeq object is also provided with both raw and cumulative sum scaling (css) normalization counts. Phyloseq is an R package to import, store, analyze, and graphically display complex OTU-clustered high-throughput phylogenetic sequencing data. The package implements advanced/flexible graphic systems and leverages many of the tools available in R for ecology and phylogenetic analysis for downstream analysis.

The report also contains the output of the taxonomic classification performed using kraken2, providing both an interactive bar plot summarizing the results of the taxonomic classification on the whole dataset both the single kraken2 and krona pie chart output file for each sample (**Fig 3, panel A**).

Finally, the last part of the report provides general statistics about the reads quality check/preprocessing and assembly metrics (calculated using QUAST tool).

### Validation and testing

MetaPhage performance was tested by analyzing a DNA virome and a shotgun metagenome data set from a published study (Liang et al. XX). The authors analyzed 20 healthy infants’ stool samples, collected at 0-4 days after birth, at 1 and 4 months of life, in order to investigate the origin of the viral population in the gut. The authors reported 2552 viral contigs, 1029 of which were classified as Bacteriophages containing more than 10 ORFs.

Our MetaPhage analysis, run with the same parameters found 4167 vOTUs (2374 of which contains more than 10 ORFs) in the DNA virome dataset and 7242 vOTUs (2562 of which contains more than 10 ORFs) in the shotgun metagenome. VirFinder, VirSorter and Phigaro gave a relatively limited list of reliable viral contigs, while Vibrant outputs a considerably larger list of candidates, on both datasets. Bacteriophages classification results, collapsed at the *Family* taxonomic rank and RPM normalized, are shown in **Figure 4**.

**Figure 4.**
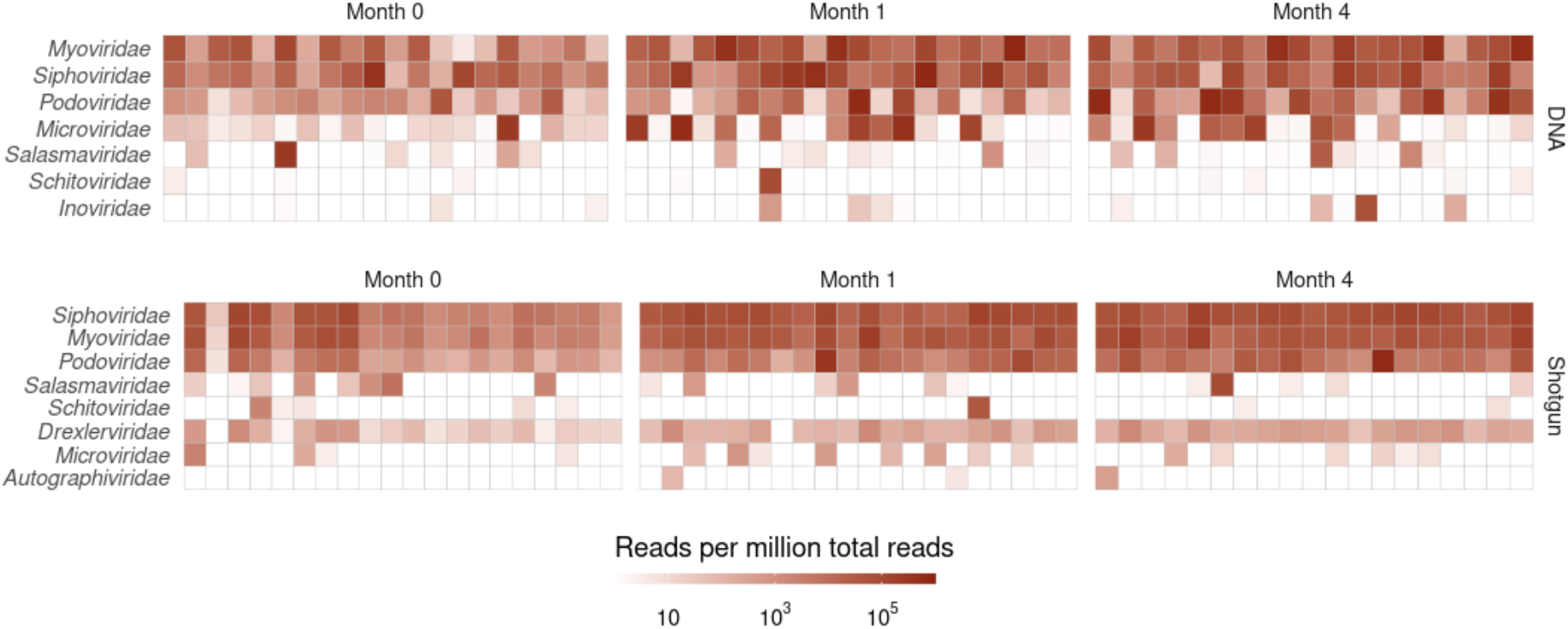
Heatmaps of the abundance of predicted viral Families across each sample in the two datasets tested.

## Discussion

MetaPhage is a modular and automated tool to detect, annotate and analyze bacteriophages present in a variety of different metagenomic datasets (DNA or RNA viromes, shotgun metagenomes) allowing the user to select and combine four phage miners for the analyses. The combination of viral contigs detected by more than one method is recommended to obtain comprehensive results, since different bacteriophage miners based on different theoretical premises can complement each other (Hayes et al., 2017). Starting from raw reads, our pipeline provides a taxonomic annotation for each assembled phagic sequence, as part of a rich interactive report made up of useful plots and tables, which can be opened on any browser and downloaded directly on the machine.

From a computational perspective, MetaPhage is a scalable pipeline, which can be run on any platform that supports Conda, Singularity or Docker. The defined dependencies between the various steps of the calculations and different intermediate level results files allow to re-run the pipeline and restart the analyses from the last concluded step, if needed. MetaPhage makes it easy to perform a complex analysis of bacteriophages in metagenomics, yielding reliable results that help understanding and visualizing the composition and structure of viral communities.

## Availability

MetaPhage is available under a GPL-3.0 license at https://github.com/MattiaPandolfoVR/MetaPhage while *graphanalyzer* is available under the same license at https://github.com/lazzarigioele/graphanalyzer. MetaPhage documentation is available online at https://mattiapandolfovr.github.io/MetaPhage.

## Contribution

MP, GL and AT developed the MetaPhage pipeline, working together with NV and EA to design MetaPhage and write the first draft; moreover, NV, AT and EA supervised the project and did the critical reading, correction and revision of the manuscript, together with the general adjustments for the metagenomic workflow.

## Funding

This work was supported by 3S_4H Project; Safe, Smart, Sustainable food for Health; ID 10065201; FESR Regione 2017; POR FESR 2014–2020 and ISCRA Italian SuperComputing Resource Allocation projects (IscraC, IsC92_MPhage) founded by CINECA. EA and AT have been supported by BBSRC Institute Strategic Programme Gut Microbes and Health BB/R012490/1 and its constituent projects BBS/E/F/000PR10353 and BBS/E/F/000PR10355. AT and NV established the collaboration thanks to a BBSRC Flexible Talent Mobility Account scheme award (BB/R506552/1). The pipeline has been tested on CLIMB-BIG-DATA computing infrastructure, funded by the UK’s Medical Research Council through grant MR/T030062/1.

## Notes

https://github.com/MattiaPandolfoVR/MetaPhage

https://github.com/lazzarigioele/graphanalyzer

### Competing Interest Statement

The authors have declared no competing interest.

